# Integrating real-time data analysis into automatic tracking of social insect behavior

**DOI:** 10.1101/2020.11.03.366195

**Authors:** Alessio Sclocco, Shirlyn Jia Yun Ong, Sai Yan Pyay Aung, Serafino Teseo

**Affiliations:** School of Biological Sciences, Nanyang Technological University, Singapore; Netherlands eScience Center, Amsterdam, The Netherlands

## Abstract

Automatic video tracking has become a standard tool for investigating the social behavior of insects. The recent integration of computer vision in tracking technologies will likely lead to fully automated behavioral pattern classification within the next few years. However, most current systems rely on offline data analysis and use computationally expensive techniques to track pre-recorded videos. To address this gap, we developed BACH (Behavior Analysis maCHine), a software that performs video tracking of insect groups in real time. BACH uses object recognition via convolutional neural networks and identifies individually tagged insects via an existing matrix code recognition algorithm. We compared the tracking performances of BACH and a human observer across a series of short videos of ants moving in a 2D arena. We found that, concerning computer vision-based ant detection only, BACH performed only slightly worse than the human observer. Contrarily, individual identification only attained human-comparable levels when ants moved relatively slow, and fell when ants walked relatively fast. This happened because BACH had a relatively low efficiency in detecting matrix codes in blurry images of ants walking at high speeds. BACH needs to undergo hardware and software adjustments to overcome its present limits. Nevertheless, our study emphasizes the possibility of, and the need for, integrating real time data analysis into the study of animal behavior. This will accelerate data generation, visualization and sharing, opening possibilities for conducting fully remote collaborative experiments.

## Introduction

Scientists apply automated pattern analysis to images of experimental animals to extrapolate measures of their behavior^1–3^. Annotating behaviors using computer algorithms, rather than employing human labor, increases experimental reliability because the performance of machines varies less than that of humans. In addition, machines easily manage to focus simultaneously on multiple individuals for long periods of time, without suffering from tiredness. This allows generating otherwise unachievable data amounts in short time windows.

In the last decade, automatic video tracking systems have emerged as ideal tools to investigate the behavior of ants^4–10^. These live in compact societies that thrive in the laboratory and easily adapt to experimental setups, performing a relatively narrow repertoire of individual actions and engaging in simple social interactions. Automatic trackers can therefore easily scrape a significant amount of information just by scanning images of their colonies. Researchers usually employ tracking systems to analyze groups of ants moving in two-dimensional arenas, tagging each individual with unique identity markers (e.g., QR or ArUco matrix codes^7,8,11,12^ or combinations of painted color dots ^4–6,13^) and using cameras to take images from top and/or bottom^14^. Many video trackers rely on inferring the position and orientation of insects via the detection of individual tags in images taken at given time intervals^7,12^. From tag positions, experimenters can then estimate a variety of informative individual attributes, such as the distance traveled between data points or the general activity levels of individuals. Combined together, tag positions, relative code orientations and distances allow reconstructing social interaction networks within groups^3,15,16^. More recent ‘next generation’ video tracking systems integrate insect identification with analyses of their shapes and movements^13^. This not only allows to automatically identify individual animals, but also to classify some of their behaviors^13,16,17^.

Although the technology underlying insect video tracking advances rapidly and significantly, most of the current systems still rely on offline data processing. This means that researchers perform tracking on previously recorded videos and analyse data only *a posteriori*. This approach allows correcting errors and employing sophisticated and costly computational strategies, or even processing earlier datasets using future knowledge. On the other hand, offline data processing does not allow for simultaneous insights into already running experiments^18^. Graphics Processing Units (GPUs), which allow accelerating computation, have instead the potential to address this gap via integrating real time data analysis^19,20^ in the traditional way of conducting tracking-based studies on insect behavior. In principle, GPU-based real time data analysis systems allow tracking experimental individuals and simultaneously analyze data while storing these for further processing, making results immediately available. Combining real-time processing with traditional experimental methods, human observers could identify individual behavioral attributes, characterize social interactions and even determine group-level emerging properties before the end of an experiment. This not only increases the potential discovery rate by accelerating data generation and condivision, but also allows for iterative adjustments of experimental parameters based on real-time results.

With the aim of advancing in such direction, we developed BACH (Behavior Analysis maCHine), a real time video tracking system based on computer vision. BACH integrates existing open source convolutional neural networks^21,22^, an object identification model^21^ and a system detecting matrix codes (ArUco^23,24^). We conducted a series of experiments to compare the performances of BACH and a human observer (HO), aiming to: 1) identify whether and how BACH made mistakes or omissions, and develop strategies to limit these; 2) determine details about the subtasks BACH and HO performed, i.e., shape-based detection and code-based identification; 3) generate data of general relevance about the performances of humans and machines in following trajectories of individually identified insects.

We found that HO executed in weeks only a fraction of the tracking work that BACH did in real time. However, HO always outperformed BACH in both ant detection and identification. Nevertheless, the degree to which the performance of HO exceeded that of BACH depended on the subtask (detection vs. identification) and on the tracking conditions (e.g., combinations of light type and ant-hosting environment). While BACH identified ants almost as efficiently as HO, its identification performance did not achieve human-like accuracy levels. HO performed better than BACH because of errors resulting from BACH’s way to detect and identify ants; in addition, while the physical limitations of our experimental prototype (camera resolution, blurred images etc.) seemingly affected the performance of BACH, they affected that of HO to a lesser extent.

Besides developing and testing another tool to conduct research on insect social interactions, this study aims to encourage researchers to integrate real time data analysis in their systems for investigating animal behavior. This may help achieve faster result generation and conduct fully remote cooperative research projects.

## Material and methods

### Experimental settings

We built a casing prototype using 5mm-thick black acrylic, aiming to host a single experimental ant colony per experiment (Figure 1a). This included a nest area (17×12×17cm) and a foraging area (17×12×18cm) connected by a 5mm-diameter tunnel, which we kept closed in this study (Figure 1b). The casing stood on a plaster of Paris base (Figure 1c), which we humidified with water prior to experiments. In the nest area, we molded three interconnected circular chambers in the plaster base, covering these with a glass lid. We painted the facing-down side of the lid with Sigmacote® (Sigma) to prevent ants from climbing onto the ceiling and walking upside down. In the foraging area, ants walked on the plaster surface, and FluonⓇ (Sigma) on the chamber walls prevented them from escaping. We positioned webcams (Logitech Brio) with the infrared filter removed in compartments above each area, and connected them using USB cables to a desktop computer (DELL Precision 7820 Tower Workstation upgraded with Dual GTX1080 GPUs and 128Gb RAM). We affixed strips of white adhesive tape to the chamber walls to minimize the proportion of dark pixels in the images, which facilitated automatic exposition and focus. Two crossed strips of tape on the plaster floor of the foraging area further facilitated the camera focus.

**Figure 1.**
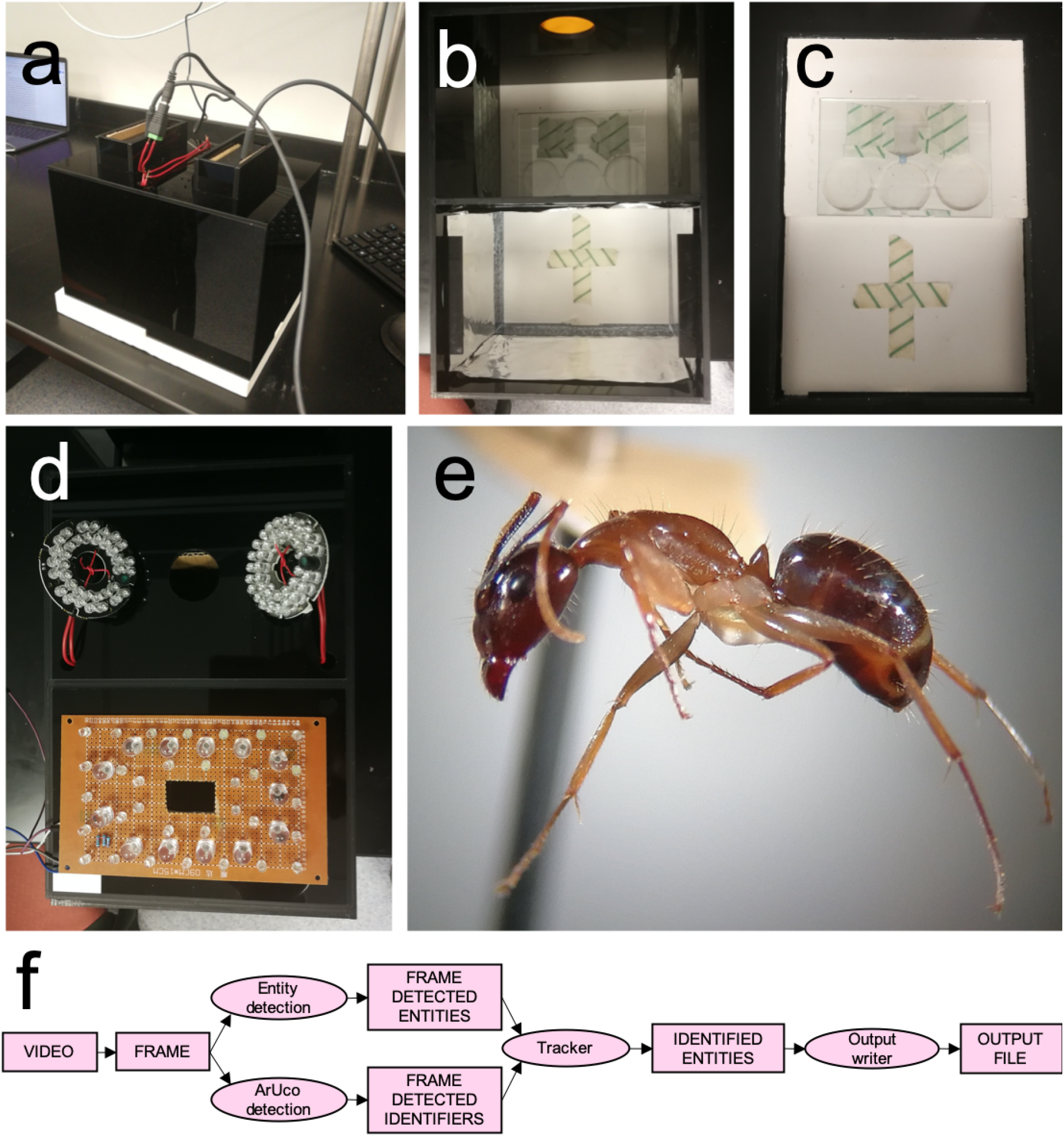
**a.** The video tracking prototype. **b.** The tracking setup interior. **c.** the plaster base with the nest area on top and the foraging area on the bottom. **d.** The illumination system: ready-made IR LED diod panel on top, veroboard with white, IR and UV led circuits on the bottom. **e.** The carpenter ants used in the experiments. **f.** A flux diagram illustrating the functioning of BACH.

### Lighting

In the nest area, two ready-made circular mini panels of 36 LED diodes, oriented at 45 degrees relative to the ceiling of the box, conveyed 850nm near infrared light (Figure 1d). Invisible to ants, near infrared light simulated the darkness of a real ant nest interior, at the same time allowing image recording via IR-sensitive webcams. In the foraging area we employed white, UV and infrared light, conveyed via three independent circuits of LED diodes (24, 4 and 13 units for white, UV and IR light, respectively) manually soldered to a stripboard (Figure 1d, video S1). We set up the lighting system to alternate UV, white and IR light in the foraging area via an Arduino microcontroller, through customizable commands.

### Ant collection and preparation

For experiments, we collected and kept in the laboratory an ant colony fragment (genus *Camponotus*, Figure 1e). We printed, hand-cut and glued 20 2×2mm 4×4-pixel ArUco markers^23,24^ (from 0 to 19, obtained via http://chev.me/arucogen/) on the abdomens of ice anesthetized ant workers, using AralditeⓇ Rapid 5 Minutes epoxy. Then, we immediately placed marked ants in an open container for recovery; finally, we either placed them in the nest chambers after a second brief ice anesthesia, or released them from the container into the foraging area through a Fluon-coated funnel placed in the webcam window above. We waited for ants to recover normal activity levels before recording videos.

### Image annotation and model training

For model training and testing, we established two sets of images including ArUco-tagged and non-tagged ants, one for the nest area and the other one for the foraging area. We extrapolated images from videos taken in settings identical or similar to those of the experimental tracking. For training and validation, we used respectively 1551 and 53 images in the nest, and 2597 and 108 in the foraging area. We manually annotated ant images using the software Yolo_mark^21^, selecting rectangular ant-including areas.

### Tracking

Using the two models resulting from the training, BACH (https://github.com/isazi/bach) tracked ants in two consecutive steps (Figure 1f). First, it tried to detect each ant using the YOLO deep neural network model^21,22^, processing each frame of the input video independently but in the same order as frames appeared in the video. Then, it added to a list all the detected entities for which the probability of looking like an ant exceeded a user-defined threshold; for each detected entity, the information stored in the list included the coordinates of the top-left and bottom-right vertices of a rectangular bounding box drawn around each ant. In the second step, BACH processed the same frame using OpenCV^25^ to detect ArUco markers. Finally, it stored each detected marker in a list containing marker ID’s and the xy coordinates of their central point (the intersection of the diagonals of the bounding box).

BACH then processed the list containing the detected entities and the one containing the detected ArUco markers. First, it compared the newly detected entities to the list of those detected in the previous frames, to measure potential overlaps between their bounding boxes. If the bounding-box of a newly detected entity did not overlap with any already known box, BACH added it to the system as a new ant with id “−1”. If the bounding-box overlapped with one or more boxes with an already known identity, BACH did not add it to the system as a newly detected entity, updating instead the position of the ant with the largest overlap with the new detection. In addition, it also marked the known ant as “seen” for the current frame. After BACH processed all entities, it engaged in a similar process for the detected ArUco markers, assigning to an entity each detected marker whose central point fell inside its bounding-box. In case the central point of a marker fell in the intersection of two or more bounding-boxes, BACH assigned it to the entity whose central point lied the closest to the marker’s central point. Each entity kept track, using an ordered list, of the count of all ArUco markers assigned to it, and used the ArUco marker with the highest count as its own ID. This ensured that the omission or incorrect detection of a marker did not affect the status of a previously detected and identified ant, allowing BACH to attribute identities to ants even when it could not see their matrix codes.

BACH deleted entities still present in the system if it could not detect them for a user-defined number of frames. In other words, in the event that a given ant detected in a frame became undetected in subsequent frames, BACH would retain its latest detection throughout a certain number of frames. We defined this as ‘ghost threshold’ because, in tracked videos, the bounding box of an ant becoming undetected remained empty in the location of its last detection. If BACH detected the ant again before reaching the ghost threshold, it kept tracking it, whereas it deleted the retained detection if it did not succeed in detecting the ant again. We set the ghost threshold to one frame (0.1 sec) for the tracking in the foraging area, and five frames (0.5 sec) for that in the nest area. We used such values because ants in the foraging area tended to move faster than in the nest, resulting in more missed detections. Finally, BACH appended all entities still present in the system to the output file of the tracker, which recorded the position of each identified entity for each frame.

### Video recording

We randomly selected four sets of around twenty workers of various sizes (one set for each of four recording days). This allowed us to account for random variation stemming from individual behavior and from the manual positioning of the matrix markers on ant bodies. We recorded videos at 10 frames/second for 25 seconds, in three different conditions: 1) foraging area exposed to infrared light; 2) foraging area exposed to visible plus UV light; 3) nest area exposed to infrared light. For each of the three conditions, we recorded ten different videos for each of four days (120 videos in total), with 10-minute pauses in between.

## Measuring system efficiency

### Frame Extraction and ant identification by the Human Observer (HO)

We randomly selected four videos per condition taken across at least three different days. Each video consisted of 250 frames labelled 0 to 249. As manual ant identification required a significant amount of repetitive work, we extracted and analysed only one in ten frames (19 frames from 60 to 240). We took frame 60 as a starting point because, after activation, the camera focus stabilized by frame 60 across all videos. HO identified ants by reading their ArUco codes and using ImageJ^26^ to compare their coordinates with those attributed by BACH (and not knowing the identities BACH gave to ants). This allowed HO to align ants with the coordinates provided by BACH and to manually annotate coordinates of ants that BACH did not detect. HO also classified each ant in terms of visibility of its ArUco marker (horizontal, tilted, invisible, Figure S1a, b, c) and blurriness (sharp, blurry, very blurry, Figure S1 d, e, f). HO also considered codes as invisible under other circumstances, including: instances in which one or more ants partially covered the body of another ant, altering its shape and/or obstructing its ArUco code (Figure S1 g); instances in which ants partially or totally climbed the foraging area walls (Figure S1 h), and as they did not contrast against the black acrylic walls of the tracking setup. In such cases, BACH could not detect them, and therefore could not locate and identify their matrix code; instances in which ants bended their abdomen forward in order to groom themselves (Figure S1 i), which altered their shape and made their ArUco codes invisible to the cameras; instances in which ants climbed on the nest glass lid ceiling and walked upside down, which hid matrix codes under their bodies (Figure S1 j).

### Verification of HO identification and manual tracking

To evaluate and compare the performances HO and BACH, we needed a completely accurate ant identification that we could use as a reference. Therefore, we further manually tracked all ants across all frames of the 12 analyzed videos. This procedure differed from the aforementioned tracking of HO, which only analyzed one frame per time without referring to other frames within the same video, and did not know the identities attributed by BACH. Contrarily, in this procedure we verified and corrected all detections, and for instances in which HO or BACH did not identify codes, we replayed videos and followed individuals until their code appeared identifiable in a preceding or following frame. In addition, we manually tracked all ‘unknown’ individuals that HO or BACH could never identify in any of the frames. Besides spotting misidentification errors, this further tracking allowed us to fill gaps between frames, for example when HO or BACH identified an individual at frame 60, did not identify it at frames 70, 80, 90, and identified it again at frame 100. Importantly, it also allowed retrieving the coordinates of all individuals across all frames, which in turn enabled us to classify attributes of the instances in which BACH and HO detected/identified them, or failed to do so.

Finally, during this manual verification, we discovered two peculiar cases of detection by BACH: in the first, which we defined as ‘double’ (Figure S1 k), BACH detected two ants as a single one; in the second, that we defined as ‘double plus one’, it first detected two ants as a single one, and then detected correctly one of the ants while still detecting both ants as a single one (Figure S1 l). In our manual tracking, we treated double detections as equivalent to detecting only one of the ants, and “double plus one detection” as equivalent to detecting both ants.

### Ant walking speed calculations

To have an idea of the walking speed of each experimental individual, we used as a proxy the distance between the xy coordinates of the same ant in two consecutive frames. This corresponded to the average walking speed of the individual during the second preceding each analyzed frame. As a consequence of this, we could not calculate the ant speed at the first analyzed frame of each video (frame 60). Therefore, we removed frame 60 in all analyses taking into account the ant walking speed.

BACH and HO retrieved the xy coordinates of ants by annotating their central point of its shape within the picture. HO did this by eye with the support of ImageJ for reading individuals’ coordinates; for BACH, an individual coordinates corresponded to the center of a bounding box including each ant shape in a given frame. Therefore, if individual positions varied slightly across two consecutive frames (e.g., an ant moved a leg without changing location), BACH’s bounding boxes changed in order to fit the new shape. This ultimately resulted in a shift of the coordinates of the bounding box’s center, which created a small, artifactual displacement the ant did not actually make. To avoid including this artifactual variation, we visually analyzed a subsample of ant displacements with the corresponding speed measures, and decided to round to zero all speed values lower than 0.75 mm/sec.

### Statistical analyses

For statistics on the number of ant detections, we used R^27^ to conduct Pearson’s chi square tests. We proceeded this way, which implied considering each video frame as independent from the previous one, because BACH detected ants within each frame independently from their position in the previous frame.

For analyzing identification data, we needed to consider interactive factors and/or used random intercepts that allowed fitting repeated measure designs on binomial or count data. Therefore, we used the R package lme4^28^ to implement Generalized Linear Mixed Models (GLMMs). For most models, we included individual identity nested in frame number nested in video as a random factor (in the R package lme4: 1|video/frame/id). We used this structure because BACH identified individuals also based on their position and identity in the previous frames, while videos involved different sets of individuals.

To test the significance of interactive effects in GLMMs, we compared models including both the interaction and the single factors with corresponding models only including single factors; similarly, to test the significance of single factors in models without interactions, we compared the model including the factor with the corresponding model only including the intercept. For such comparisons, we used the R function ‘anova’ with ‘Chisq’ test specification. To test specific *post hoc* contrasts, we used the R package emmeans^29^ (with Tukey HSD test and automatically adjusted p values) when the factors of interest had more than two levels. When factors had only two levels, we instead referred to the contrasts appearing in the output of the R function ‘summary’ applied to each model. If the residual deviance considerably exceeded the residual degrees of freedom, we corrected GLMs and GLMMs for overdispersion by including each observation as an additional random factor. If models failed to converge or resulted in singular fits, we used optimizers (from the optimx R package^30,31^). We provide scripts for statistical tests in the supplementary material.

## Results

### Ant detection

HO detected ants significantly better than BACH, overall (Pearson’s chi-squared test, χ^2^=725.68, df=1, p<0.001) and in each of the three tracking conditions (Pearson’s chi-squared tests, nest: χ^2^=212.17, df=1, p<0.001; foraging area under visible light: χ^2^=227.28, df=1, p<0.001; foraging area under infrared light: χ^2^=288.64, df=1, p<0.001, Figure 2a). Considering all conditions together, HO detected ant shapes in all instances, while BACH did so 83.8% of times. As HO never failed to detect ants, its accuracy did not vary across conditions. On the other hand, BACH detected 86.6% of ant shapes within the nest, 84.7% in the foraging area with visible light and 79.3% in the foraging area with infrared light, with a significant condition-specific variation in detection efficiency (Pearson’s chi-squared test, χ^2^=27.945, df=2, p<0.001, Figure 2a).

**Figure 2.**
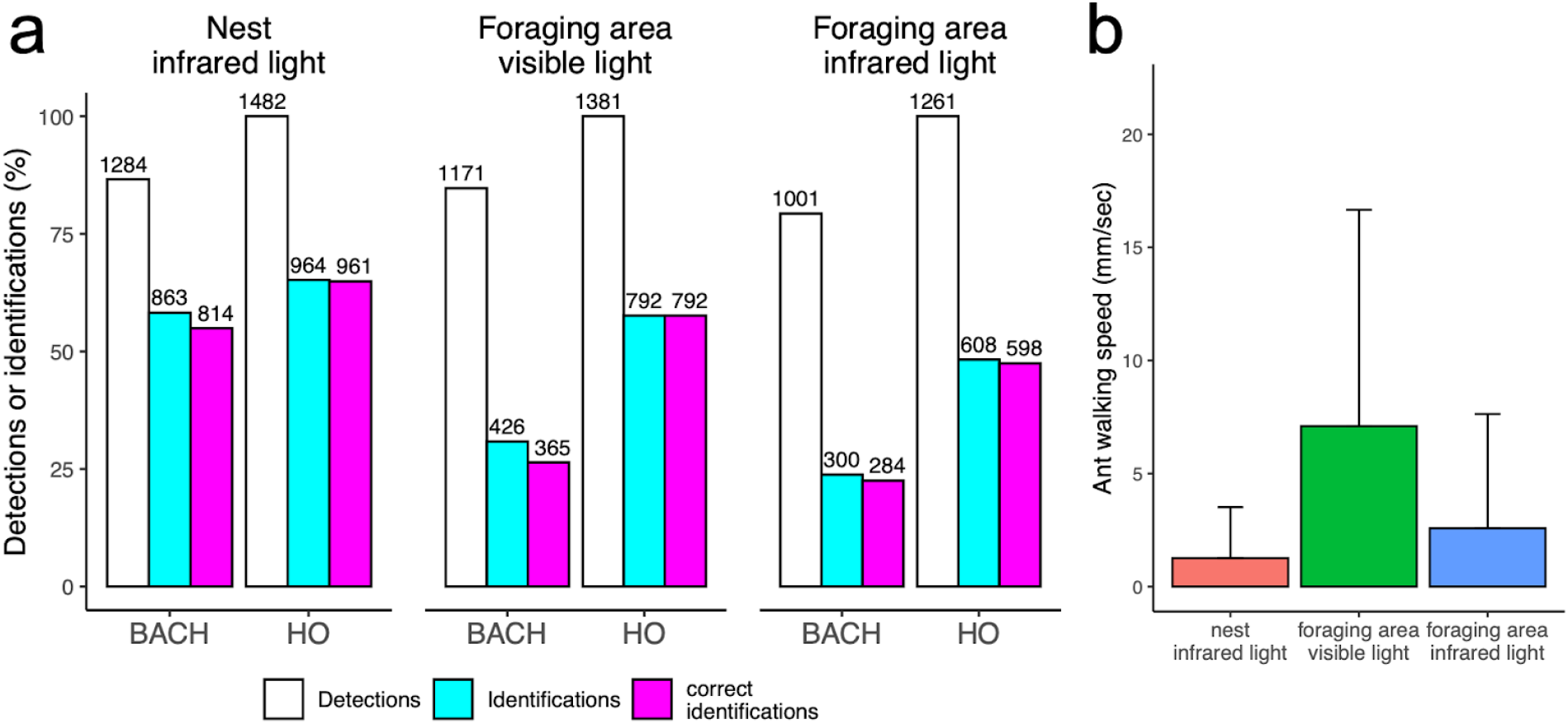
**a.** Number of detections, identifications and correct identifications for BACH and HO across conditions. b. Walking speed of ants across conditions (mean + sd).

The walking speed of ants varied significantly across conditions (GLMM, χ^2^=12.699, df=2, p<0.01, Figure 2b). Ants walked the fastest in the foraging area under visible light (6.71±9.4 mm/sec), significantly faster than the average 2.44±4.9 mm/sec of ants in the foraging area under infrared light (estimate=2.225, z=3.484, p<0.01) and than the average 1.19±2.2 mm/sec of ants inside the nest (estimate=2.888, z=4.521, p<0.001). In infrared light conditions, ant speed did not vary between nest and foraging area (GLMM, estimate=0.663, z=1.032, p=0.55). BACH-detected ants walked faster than undetected ants, in general (GLMM, estimate=0.07378, z-value=6.653, p<0.001) and in each tracking condition (GLMMs; nest: estimate=0.17, z-value=3.492, p<0.001; foraging area under infrared light: estimate=0.11, z-value=4.288, p<0.001; foraging area under visible light: estimate=0.05, z-value=4.665, p<0.001, Figure 3a). HO detected ants in all instances, therefore ant walking speed did not affect its performance.

**FIgure 3.**
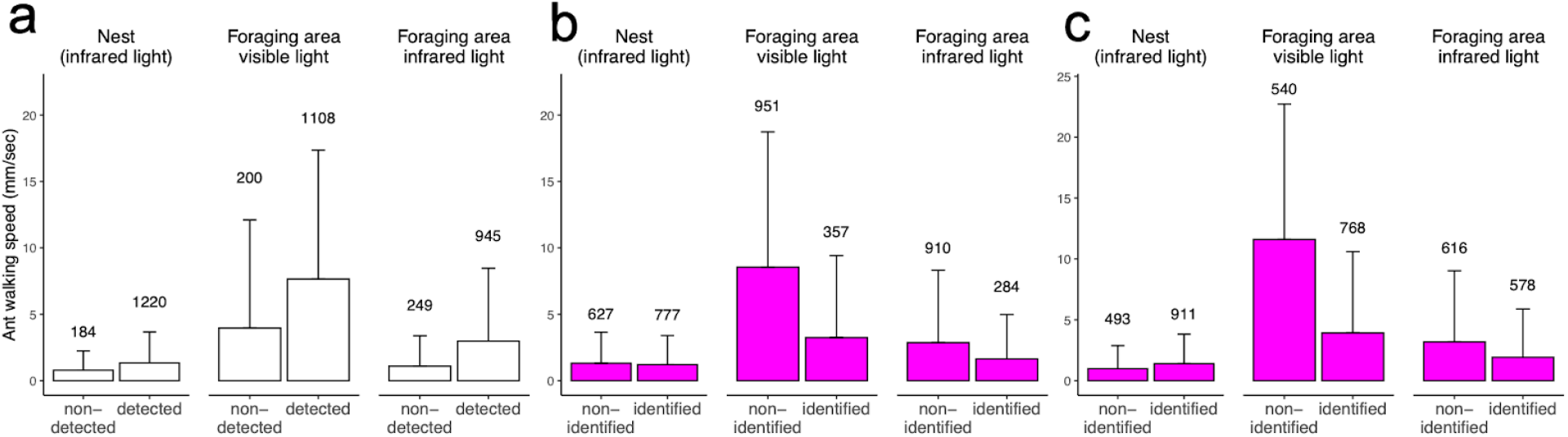
**a.** Speed of BACH-detected and undetected ants in different conditions.. **b.** Speed of BACH-identified and non-identified ants. **c.** Speed of HO-identified and non-identified ants.

**Figure 4.**
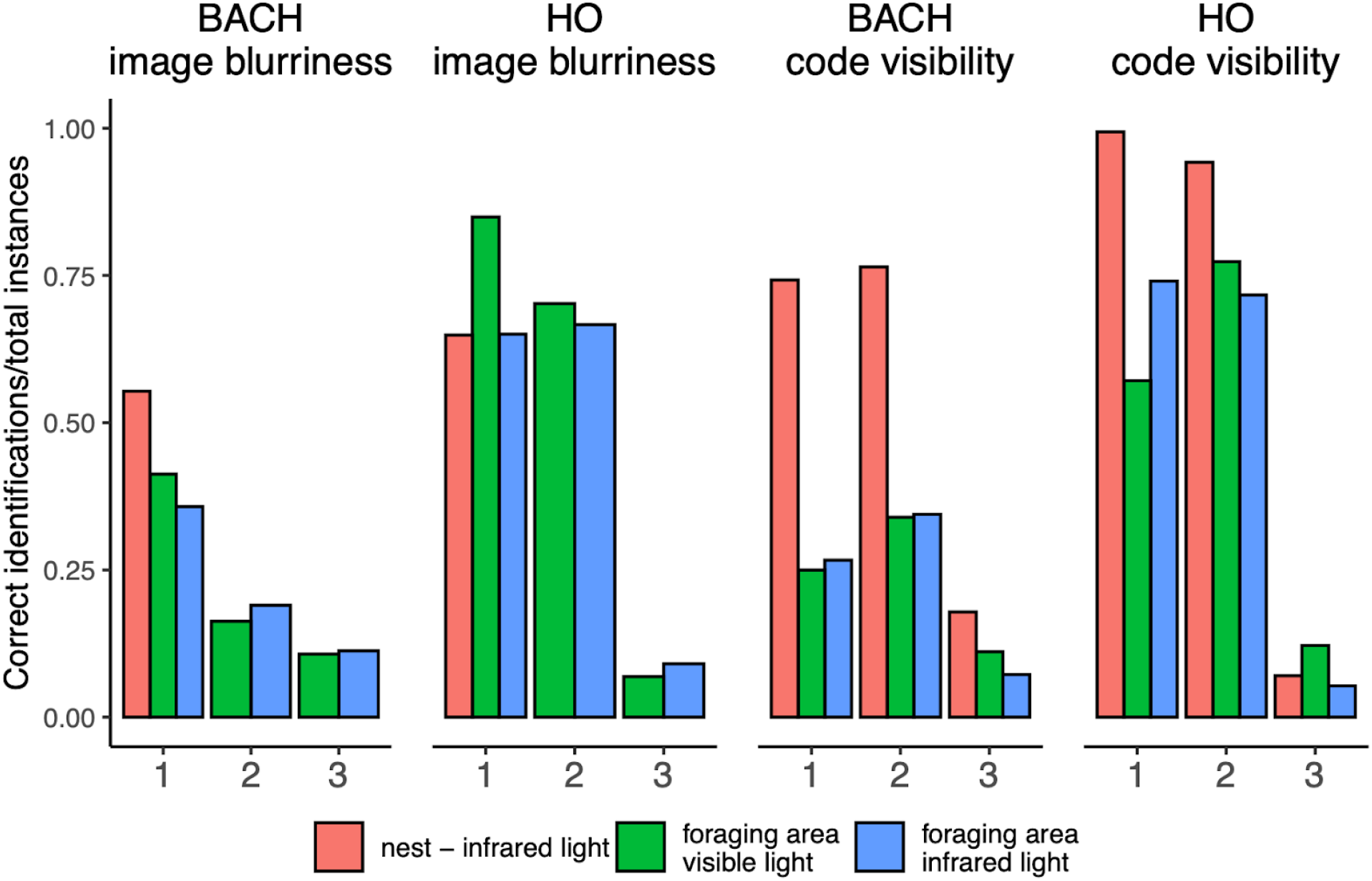
Proportion of identifications by BACH and HO across different classes of blurriness and code visibility. Numbers correspond to the different classes (sharp, blurry, very blurry for image blurriness, horizontal, tilted, invisible for code visibility).

Sometimes BACH detected two ants as a single one, at a rate that varied significantly across conditions (Pearson’s chi-squared test, χ^2^=11.31, df=2, p<0.01). BACH produced 72 of such “double detections” inside the nest (4.8% of total nest detections), 48 in the foraging area with visible light (3.4%), and 79 in the foraging area with infrared light (6.2%). “Double plus one” detections occurred instead only once in the nest (0.13%), 6 times in the foraging area under visible light (0.36%), and 9 times in the foraging area under infrared light (0.72%), also varying significantly across conditions (Pearson’s chi-squared test, χ^2^=7.4785, df=2, p=0.023). In two of the videos taken in the nest area, we did not apply Sigmacote^®^ on the facing down side of the glass covering the nest chambers, which resulted in ants walking on the nest glass ceiling. Of 238 instances of ants walking on the glass ceiling (31.3% of total detections in the two videos), BACH failed to detect ants 44 times (18.4%). Finally, sometimes ants partially or totally climbed the foraging area walls. In some of such cases, BACH could not detect them. This occurred 98 times in the foraging area in visible light and 78 times in the foraging area in infrared light (respectively, 7% and 6.1% of total detections in each condition).

### Ant identification

HO identified ants better than BACH across all conditions (GLMM, nest: estimate=0.88, z=8.61, p<0.001; foraging area in infrared light: estimate=1.89, z=17.16, p<0.001; foraging area in visible light: estimate=1.95, z=20.38, p<0.001, Figure 2a). Ant walking speed, ant image blurriness and code visibility interactively affected both BACH’s and HO’s ant identification (anovas on GLMMs, respectively: χ^2^=122.81, df= 12, p<0.001; χ^2^=954.18, df= 12, p<0.001). However, non-identified ants generally walked faster than identified ants (BACH: χ^2^=139.25, df=1, p<0.001, Figure 3b; HO: χ^2^=237.83, df=1, p<0.001, Figure 3c), tending to appear blurry in many video frames. Accordingly, ant walking speed varied significantly for different code blurriness classes (namely: ‘sharp’, ‘blurry’, ‘very blurry’; anova on GLMMs, ‘Chisq’ test, χ^2^=303.61, df=2, p<0.001). In particular, sharp ants walked at lower speeds compared to blurry and very blurry ants (*post hoc* contrasts, respectively: estimate: −1.23, z-ratio=−7.814, p<0.001; estimate: −2.72, z-ratio=−17.494, p<0.001), and blurry ants walked at lower speeds compared to very blurry ants (estimate:−1.49, z-ratio=−8.264; p<0.001). This corresponded to different walking speeds in the foraging area (under visible light: sharp: 3.6±6.99 mm/sec, slightly blurry: 6.18±8.99 mm/sec, very blurry: 6.57±9.09 mm/sec; under infrared light: sharp: 2.09±4.28 mm/sec, slightly blurry: 3.08±5.68 mm/sec, very blurry: 5.66±8.86 mm/sec). We only found sharp ant images in videos taken inside the nest, where ants generally walked at a lower speed (1.19±2.2 mm/sec) compared to the other conditions. We concluded that BACH may have had difficulties in identifying fast-walking ants because of their blurry codes. We therefore dropped speed in favor of image blurriness in further analyses.

We found an interactive effect of image blurriness and code visibility on identification accuracy for both BACH and HO, overall (anovas on GLMMs, respectively: χ^2^=111.19, df= 4, p<0.001; χ^2^=1102.3, df= 4, p<0.001) and in the foraging area, both under infrared light (BACH: χ^2^=71.816, df= 4, p<0.001; HO: χ^2^=133.98, df= 4, p<0.001) and visible light (BACH: χ^2^=23.338, df= 4, p<0.001; HO: χ^2^=401.59, df= 4, p<0.001, significant contrasts for each condition in Table S1). As in the nest area we only encountered sharp ant images, we only tested the effect of code visibility, which we found significant for both BACH (anova on GLMMs, χ^2^=460.87, df= 2, p<0.001) and HO (anova on GLMMs, χ^2^=1333, df= 2, p<0.001). In general, and as expected, BACH and HO tended to best identify ants in sharp images and when they could see their codes.

Both HO and BACH occasionally attributed wrong identities to ants (Figure 2a). For all conditions pooled, HO identified ants in 57.46% of all instances, accurately doing so 57.15% of times. BACH identified ants in 38.53% of instances, with 35.47% of correct identifications. In the nest area, HO identifications reached 65.18% of the total, with 64.9% of correct identifications; in the foraging area under visible light, HO identified ants in 57.6% of instances, never making mistakes; finally, in the foraging area under infrared light, it identified codes 48.25% of times, with 47.46% accuracy. In the nest area, BACH identified codes 58.23% of times, with 54.92% correct identifications; in the foraging area under visible light, it identified codes the 30.84% of times, with 26.43% accuracy; in the foraging area under infrared light, identifications dropped at 23.79% of the total, and correct identifications at 22.52%. The ratio of correct to total identifications of HO significantly exceeded that of BACH across conditions (nest: Pearson χ^2^ test, χ^2^=47.536, df= 1, p<0.001; foraging area in visible light: Pearson χ^2^ test, χ^2^=119.97, df= 1, p<0.001; foraging area under infrared light: Pearson χ^2^ test, χ^2^=9.8263, df= 1, p<0.01). This indicated that, when identifying codes, BACH made more mistakes than HO. BACH attributed incorrect identities in 165 of 1628 identification instances (10.13%). These mistakes resulted from 27 identity exchange events between ants located in close proximity (7.46±3.52mm). In such occasions, BACH did not disentangle the shapes of different individuals, for example detecting one ant shape and part of the other ant shape as a single entity. Even if such mistaken detections lasted only for one or few frames, they ultimately resulted in identities to shift from an individual to another. BACH kept attributing the shifted identity through a varying number of frames (6±5.15), typically until it managed to read again the ArUco code of the identity-shifted individual. Identification errors of HO consisted instead in wrong readings of the ArUco codes.

## Discussion

In this study, we developed and tested BACH, a real time video tracking system for insects based on computer vision and matrix code recognition. We then compared the performances of BACH and a human observer (HO), evaluating their efficiency in 1) detecting ant shapes and 2) identifying individuals via integrating the detected shapes with matrix codes. We found that, although HO employed weeks to track a fraction of the images BACH tracked in real time, it always qualitatively outperformed BACH, making significantly less mistakes and omissions. However, this varied depending on the subtask and the tracking conditions.

For ant detection only, BACH generally performed only slightly (although significantly) worse than HO across all conditions. Interestingly, BACH-detected ants walked generally faster compared to their undetected counterparts. This suggests that, with the detection models it employed, BACH achieved a better efficiency for moving rather than non-moving entities. This probably resulted from the higher shape variation of walking compared to non-moving ants, which increased the probabilities for BACH to encounter familiar shapes it already knew from the training phase. BACH also had a fairly high efficiency when detecting ants walking relatively fast, even though these appeared blurry in the frames. This means that BACH detected unfocused ant shapes nearly as efficiently as focused shapes, probably because the training image set also included blurry images of ants.

BACH had a low detection efficiency in a few cases, for example when ants partially climbed on the black walls of the setup casing, which made them indistinguishable from the background. In future studies, we could solve this problem by using fair rather than black acrylic, and/or by including more images of ants attempting to climb the setup’s walls in our training sets. BACH’s detection performance also fell when multiple ants engaged in clusters (‘double’ and ‘double plus one’ detections), because telling apart overlapping ant shapes became difficult. Although we still need to implement a solution for this in BACH, a recently developed tracking system (AnTrax^13,26^) has elegantly worked around this issue. This system, optimized for cluster-formic Clonal Rider Ants, retains the identities of individuals when they join a cluster and become indistinguishable from other ants. When such individuals leave the cluster, AnTrax retrieves their identity. This process allows maintaining a coherent flow of information even without knowing individual positions within the cluster at all times. In future versions of BACH, we aim to integrate a similar process that will use individual histories to retain their identities while they engage in clusters. In principle, as BACH runs modules in parallel, this will not compromise its real-time performance.

Contrary to ant detection, BACH’s identification efficiency based on matrix codes varied dramatically across tracking conditions. While BACH identified ants only slightly less effectively than HO in the nest area, its performance fell to about half of that of HO in the foraging area. This probably resulted from ants walking faster in the foraging area compared to the nest, which most likely occurred because the UV and visible light, as well as the absence of shelter, disturbed the predominantly nocturnal *Camponotus* ants. Our manual tracking and verification revealed that higher walking speed produced more instances of blurry images and therefore blurred matrix codes, which BACH often failed to identify. Accordingly, and contrary to what we found for ant detection, BACH-identified ants walked slower than non-identified ants, confirming that BACH made most identification mistakes for ants walking relatively fast. To solve this issue, we should invest future efforts in facilitating matrix code identification. From a hardware perspective, we will employ cameras with higher resolution and frame rate, which would reduce the occurrence of blurry images and thus maximize code readability. In addition, in our videos, the foraging area surface took around half of the frame, and the nest area only one tenth of it, leaving a large portion unused. In the future, we could easily cut off such extra space by reducing the distance between the camera and the tracking surface, which would result in a better resolution and therefore higher chances for BACH to correctly read matrix codes. On the software side, we will increase ant identification efficiency by improving the way BACH matches new detections to the current state of the system. For example, we could match new unknown ant detections with previously detected entities that have become unidentified, merging their status based on numbers and identities of ants tracked in previous frames. This would likely limit not only the instances in which BACH fails to detect ants, but also those in which it mistakenly identifies matrix codes.

Similar to BACH, other recent tracking systems for insects integrate convolutional networks and individual tags. However, these differ from BACH in their functioning and scopes. For example, AnTrax^13^ extracts ant-containing image portions within pre-recorded videos linking them across frames and reconstructing trajectories for identifying individuals. Contrarily, BACH processes video frames in real-time, detecting individual shapes and their IDs, only keeping track of their position. While AnTrax bases its accuracy on more complex and computationally heavier techniques currently less suitable for real-time processing, BACH works in real time but extrapolates less information and makes more mistakes. In addition, AnTrax can work in concert with JAABA^16^ to automatically classify ant behaviors, whereas BACH does not include similar features at its current state. In the future, however, we aim to combine the Python interfaces of DeepLabCut^3^ or DeepPoseKit^15^ with Yolo, integrating BACH with pose and behavior estimation. Finally, AnTrax identifies ants via color combinations, which provides advantages but also disadvantages compared to the matrix code system supported by BACH. Only ants with sufficiently large bodies can bear readable matrix codes, and yet these restrict their movements, probably affecting their behavior; sometimes matrix codes detach from the ant bodies, disrupting the information flow within experiments^13^. On the other hand, color-based identification relies on experiments conducted in visible light, which does not affect virtually blind Clonal Raider Ants but does not suit experimental work on non-blind, lucifugous or nocturnal species living in dark nests. As shown in our experiments on *Camponotus*, tracking colonies of such ants in visible light may alter their natural behavior.

Other automatic trackers used to study insect societies integrate computer vision and/or matrix code identification to different extents, working mainly offline. For example, a recent system for tracking honeybee colonies^17,32^ not only uses matrix codes for individual identification, but also allows automatically and simultaneously detecting multiple mouth to mouth food exchanges (trophallaxis). The system identifies these episodes based on the relative position and orientation of matrix codes, confirming such identifications via custom computer vision algorithms. Contrarily, AntVis^33^ employs computer vision to visualize fluxes of ants moving in the same direction, without implementing individual ant identification over extended periods of time. Conceived for observation in natural conditions, AntVis identifies individual ants by accurately tracking their trajectories within the video frame, without relying on individual matrix code tags^34^. This means that it identifies individuals as long as they continuously appear within the video frame, considering them as new whenever they exit and re-enter the frame.

Compared to other tracking systems, BACH has two main advantages. First, it remembers the identities of individuals even when their identity codes become invisible, and can successfully do so as long as it continuously detects their shape. Secondly, and most importantly, BACH generates data in real time through relatively light computational processes, which constitutes its main novelty and strength. Although BACH needs to undergo upgrading from several perspectives, we developed it with the goal of emphasizing the possibility for real time data to ameliorate animal behavior research that relies on video tracking. Real time data shortens the delay between observation and data generation to virtually non-existent, enabling researchers to alter experimental conditions based on current results. In addition, results become immediately available for sharing, which favors collaborative projects and accelerates the discovery rate. In a conceivable design that we aim to achieve in the near future, the current BACH main module will produce positional and behavioral data in real-time from video sources, while other modules will concurrently process the generated data and produce structured information. As an example, we will manage to observe, in real-time, the development of a colony’s social graph from within BACH.

Real time data analysis increasingly benefits cooperative research in multiple fields, including radio astronomy^20,35^ and physics^36,37^. Accordingly, we conceived BACH with the goal of attracting attention on how this could facilitate animal behavior research. For example, real time tracking systems like BACH may help animal scientists establish and maintain collaborations during periods of mobility restrictions. Or, in the foreseeable future, could allow conducting animal behavior experiments entirely online, with a research group broadcasting videos of animals and other groups remotely tracking and analyzing them, cooperatively and in real time.

## Supporting information

stats script

## Acknowledgements

This work was supported by a Presidential Postdoctoral Fellowship (M408080000) from Nanyang Technological University (NTU) to ST.

---

**Figure S1.**
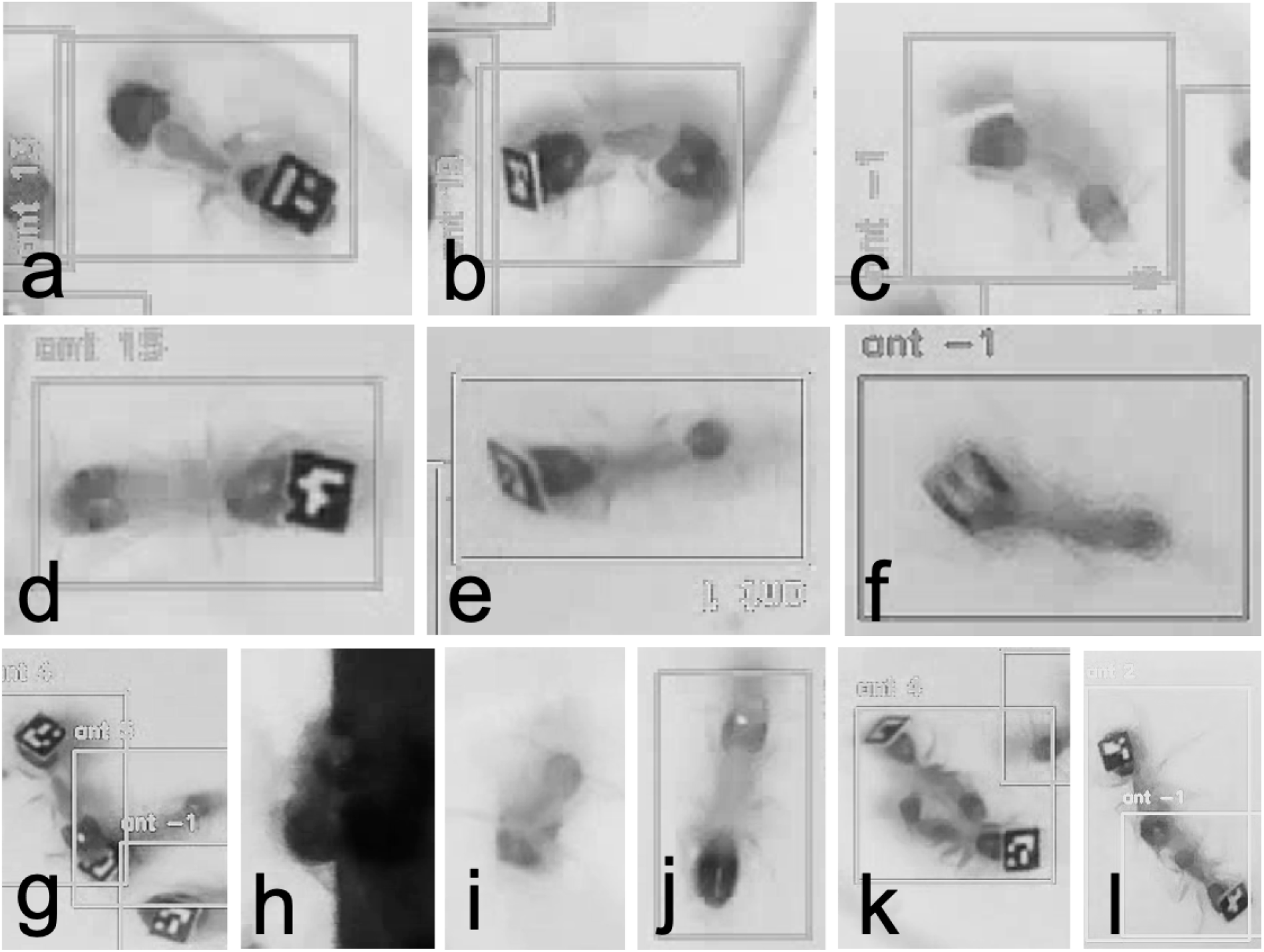
**a.- c.** Classes of code visibility (horizontal, tilted, invisible). **d.-f.** Classes of image blurriness (sharp, blurry, very blurry). **g.-j.** Examples of instances in which codes became invisible: **g.** an ant covering another ant’s code; **h.** an ant trying to climb the tracking setup wall; **i.** an ant grooming her gaster; **j.** an ant walking on the lower side of the glass lid covering the nest chambers. **k.** ‘Double detection’. **l.** ‘Double plus one detection’.

## References

1. Noldus, L. P. J. J., Lucas P J, Spink A J & Tegelenbosch, R. A. J. Computerised video tracking, movement analysis and behaviour recognition in insects. Computers and Electronics in Agriculture vol. 35 201–227 (2002).

2. Pérez-Escudero, A., Vicente-Page, J., Hinz, R. C., Arganda, S. & de Polavieja, G. G. idTracker: tracking individuals in a group by automatic identification of unmarked animals. Nature Methods vol. 11 743–748 (2014).

3. Mathis, A. et al. DeepLabCut: markerless pose estimation of user-defined body parts with deep learning. Nat. Neurosci. 21 1281–1289 (2018).

4. Ulrich, Y., Burns, D., Libbrecht, R. & Kronauer, D. J. C. Ant larvae regulate worker foraging behavior and ovarian activity in a dose-dependent manner. Behavioral Ecology and Sociobiology vol. 70 1011–1018 (2016).

5. Ulrich, Y., Saragosti, J., Tokita, C. K., Tarnita C E & Kronauer, D. J. C. Fitness benefits and emergent division of labour at the onset of group living. Nature 560 635–638 (2018).

6. Ulrich, Y. et al. Emergent behavioral organization in heterogeneous groups of a social insect. doi:10.1101/2020.03.05.963207.

7. Stroeymeyt, N. et al. Social network plasticity decreases disease transmission in a eusocial insect. Science vol. 362 941–945 (2018).

8. Richardson, T. O., Liechti, J. I., Stroeymeyt, N., Bonhoeffer, S. & Keller, L. Short-term activity cycles impede information transmission in ant colonies. PLoS Comput. Biol. 13, e1005527 (2017).

9. Heyman, Y., Shental, N., Brandis, A., Hefetz, A. & Feinerman, O. Ants regulate colony spatial organization using multiple chemical road-signs. Nat. Commun. 8 15414 (2017).

10. Greenwald, E. E., Baltiansky, L. & Feinerman, O. Individual crop loads provide local control for collective food intake in ant colonies. Elife 7, (2018).

11. Quque, M. et al. Hierarchical networks of food exchange in the black garden ant Lasius niger. Insect Sci. (2020) doi:10.1111/1744-7917.12792.

12. Mersch, D. P., Crespi, A. & Keller, L. Tracking individuals shows spatial fidelity is a key regulator of ant social organization. Science 340 1090–1093 (2013).

13. Saragosti DJC, A. G. anTraX: high throughput video tracking of color-tagged insects. BioRxiv (2020).

14. Greenwald, E., Segre, E. & Feinerman, O. Ant trophallactic networks: simultaneous measurement of interaction patterns and food dissemination. Sci. Rep. 5 12496 (2015).

15. Graving, J. M. et al. DeepPoseKit, a software toolkit for fast and robust animal pose estimation using deep learning. Elife 8, (2019).

16. Kabra, M., Robie, A. A., Rivera-Alba, M., Branson, S. & Branson, K. JAABA: interactive machine learning for automatic annotation of animal behavior. Nat. Methods 10 64–67 (2013).

17. Gernat, T. et al. Automated monitoring of behavior reveals bursty interaction patterns and rapid spreading dynamics in honeybee social networks. Proc. Natl. Acad. Sci. U. S. A. 115 1433–1438 (2018).

18. Sclocco, A. & Teseo, S. Microbial associates and social behavior in ants. Artificial Life and Robotics (2020) doi:10.1007/s10015-020-00645-z.

19. Halyo, V. V. et al. First evaluation of the CPU, GPGPU and MIC architectures for real time particle tracking based on Hough transform at the LHC. Journal of Instrumentation vol. 9 P04005–P04005 (2014).

20. Sclocco, A., van Leeuwen, J., Bal H E & van Nieuwpoort, R. V. Real-time dedispersion for fast radio transient surveys, using auto tuning on many-core accelerators. Astronomy and Computing vol. 14 1–7 (2016).

21. Redmon, J., Divvala, S., Girshick, R. & Farhadi, A. You Only Look Once: Unified, Real-Time Object Detection. 2016 IEEE Conference on Computer Vision and Pattern Recognition (CVPR) (2016) doi:10.1109/cvpr.2016.91.

22. Redmon, J. Darknet: Open Source Neural Networks in C. http://pjreddie.com/darknet/ (2013-2016).

23. Garrido-Jurado, S., Muñoz-Salinas, R., Madrid-Cuevas F J & Medina-Carnicer, R. Generation of fiducial marker dictionaries using Mixed Integer Linear Programming. Pattern Recognition vol. 51 481–491 (2016).

24. Romero-Ramirez, F. J., Muñoz-Salinas, R. & Medina-Carnicer, R. Speeded up detection of squared fiducial markers. Image and Vision Computing vol. 76 38–47 (2018).

25. Bradski, G. The OpenCV Library. Dr. Dobb’s Journal of Software Tools (2000).

26. Schneider, C. A., Rasband W S & Eliceiri, K. W. NIH Image to ImageJ: 25 years of image analysis. Nat. Methods 9 671–675 (2012).

27. R Core Team. R: A language and environment for statistical computing. R Foundation for Statistical Computing, Vienna, Austria. (2020).

28. Bates, D., Mächler, M., Bolker, B. & Walker, S. Fitting Linear Mixed-Effects Models Usinglme4. Journal of Statistical Software vol. 67 (2015).

29. Lenth R, Buerkner P, Herve M, Love J, Riebl H, Singmann H. emmeans: Estimated Marginal Means, aka Least-Squares Means. (2020).

30. Nash J C & Varadhan, R. Unifying Optimization Algorithms to Aid Software System Users:optimxforR. Journal of Statistical Software vol. 43 (2011).

31. Nash, J. C. On Best Practice Optimization Methods inR. Journal of Statistical Software vol. 60 (2014).

32. Geffre, A. C. et al. Honey bee virus causes context-dependent changes in host social behavior. Proc. Natl. Acad. Sci. U. S. A. 117 10406–10413 (2020).

33. Hu, T. et al. AntVis: A web-based visual analytics tool for exploring ant movement data. Visual Informatics vol. 4 58–70 (2020).

34. Imirzian, N. et al. Automated tracking and analysis of ant trajectories shows variation in forager exploration. Sci. Rep. 9 13246 (2019).

35. AMBER: A real-time pipeline for the detection of single pulse astronomical transients. SoftwareX 12 100549 (2020).

36. Aaij, R. et al. Design and performance of the LHCb trigger and full real-time reconstruction in Run 2 of the LHC. J. Instrum. 14, P04013 (2019).

37. Solli, D. R., Chou, J. & Jalali, B. Amplified wavelength–time transformation for real-time spectroscopy. Nat. Photonics 2 48–51 (2007).

