## Supplementary material for "Integrating real-time data analysis into automatic tracking of social insect behavior": stats script: BACH_stats.R

```
library(lme4)
library(optimx)
library(emmeans)

data1=read.table(file.choose(),header=TRUE,sep=",", dec=".")  


names(data1)
head(data1)
summary(data)
str(data1)
attach(data1)

data1$condition = as.factor(data1$condition)
data1$track_id = as.factor(data1$track_id)
data1$clarity1=as.factor(data1$clarity1)
data1$distadj = ifelse(data1$distmm_new < 0.75, 0, data1$distmm_new) ### this makes distances <0.75mm to become zero, which helps reducing false variation due to variation in ant shape detection
data1$distadj100 = round(data1$distadj, digits = 2)*100 ### this is in order to build a model with poisson distribution when distance is used as a response variable
data1$distadj10 = round(data1$distadj, digits = 1)*10
data1$frame1=(data1$frame-50)/10
data1$bach_accuracy = data1$computer_accuracy
data1$computer_accuracy[is.na(data1$computer_accuracy)] <- 0
data1$HO_accuracy = data1$human_accuracy
data1$human_accuracy[is.na(data1$human_accuracy)] <- 0

### function to calculate overdispersion
overdisp_fun <- function(model) {
  rdf <- df.residual(model)
  rp <- residuals(model,type="pearson")
  Pearson.chisq <- sum(rp^2)
  prat <- Pearson.chisq/rdf
  pval <- pchisq(Pearson.chisq, df=rdf, lower.tail=FALSE)
  c(chisq=Pearson.chisq,ratio=prat,rdf=rdf,p=pval)
}

data2= subset(data1, frame!='60') ### I exclude the first frame because speed is always zero, because our speed is referred to the previous frame, and 60 is the first frame. Keeping that frame may create a bias, I think
detach(data1)
attach(data2)
obs_effect = 1:nrow(data2)


### start with chisquare tests

total = matrix(c(4124,3457,0,667), ncol=2)
chisq.test(total,correct=F)

nest = matrix(c(1482,1284,0,198), ncol=2)
chisq.test(nest,correct=F)

FV = matrix(c(1381,1171,0,210),ncol=2)
chisq.test(FV,correct=F)

FI = matrix(c(1261,0,1002,259),ncol=2)
chisq.test(FI,correct=F)

system = matrix(c(1284,1171,1001,198,210,260),ncol=2)
chisq.test(system,correct=F)

human = matrix(c(1482,1381,1261,0,0,0),ncol=2)
chisq.test(human,correct=F)


### this is for quantifying and comparing ant speed across treatments

mean(subset(data1, condition=='nestinfrared')$distadj)
sd(subset(data1, condition=='nestinfrared')$distadj)

mean(subset(data1, condition=='foraginginfrared')$distadj)
sd(subset(data1, condition=='foraginginfrared')$distadj)

mean(subset(data1, condition=='foragingvisible')$distadj)
sd(subset(data1, condition=='foragingvisible')$distadj)

### to compare ant walking speed across conditions

ma=glmer(distadj10 ~ condition + (1|video) + (1|obs_effect), family='poisson', data=data2)
summary(ma)

mb=glmer(distadj10 ~  (1|video) + (1|obs_effect), family='poisson', data=data2)
summary(mb)

anova(ma,mb, test='Chisq')

emm = emmeans(ma, ~ condition)
pairs(emm)

#this is for effect of speed and condition on ant shape detection by BACH (how ant speed varies between detected and undetected ants)

mdet=glmer(computer_shape ~ distadj + (1|video), family='binomial', data=data2)
summary(mdet)

mdetnest=glmer(computer_shape ~ distadj + (1|video), family='binomial', data=subset(data2, condition=='nestinfrared'))
summary(mdetnest)

mdetFV=glmer(computer_shape ~ distadj + (1|video), family='binomial', data=subset(data2, condition=='foragingvisible'))
summary(mdetFV)

mdetFI=glmer(computer_shape ~ distadj + (1|video), family='binomial', data=subset(data2, condition=='foraginginfrared'))
summary(mdetFI)

### now for HO

hmdet=glmer(human_shape ~ distadj + (1|video), family='binomial', data=data2)
summary(hmdet)

hmdetnest=glmer(human_shape ~ distadj + (1|video), family='binomial', data=subset(data2, condition=='nestinfrared'))
summary(hmdetnest)

hmdetFV=glmer(human_shape ~ distadj + (1|video), family='binomial', data=subset(data2, condition=='foragingvisible'))
summary(hmdetFV)

hmdetFI=glmer(human_shape ~ distadj + (1|video), family='binomial', data=subset(data2, condition=='foraginginfrared'))
summary(hmdetFI)

### double detection differences across groups (chisquare test)

double = matrix(c(1449,1333,1181,72,48,79),ncol=2)
chisq.test(double,correct=F)

### double plus one detection differences across groups (chisquare test)

doublep = matrix(c(1481,1375,1252,1,6,9),ncol=2)
chisq.test(doublep, correct=F)

### ant detection analyses stop here

#-----------------------------------------------------------------------

### analyses on ant identifications

### compares accuracy of BACH and HO across treatments, requires a dataset in long format

binomial=read.table(file.choose(),header=TRUE,sep=",", dec=".")  
binomial$track_id = as.factor(data1$track_id)
binomial$frame=as.numeric(binomial$frame)
selected=c(0:19) #creates a dummy variable to proceed with selecting only the identifiable codes
binomial2=binomial[binomial$track_id %in% selected,] # a data set with only those individuals that BACH and HO managed to identify, as there are some (those with code number>19) were never identified by BACH or HO because their codes were invisible

accbinintnested = glmer(accuracy~condition/identifier + (1|video/frame/track_id), family='binomial', data=binomial2, control = glmerControl(optimizer = "optimx", calc.derivs = FALSE,
                                                                                                                                            optCtrl = list(method = "nlminb", starttests = FALSE, kkt = FALSE)))
summary(accbinintnested)

### 3-way interactions  
# BACH

interactive=glmer(computer_accuracy~tilt1*clarity1*distadj + (1|video/frame/track_id), family=binomial, data=data2, control = glmerControl(optimizer = "optimx", calc.derivs = FALSE,
                                                                                                                                           optCtrl = list(method = "nlminb", starttests = FALSE, kkt = FALSE)))
interactive1=glmer(computer_accuracy~tilt1+clarity1+distadj + (1|video/frame/track_id), family=binomial, data=data2, control = glmerControl(optimizer = "optimx", calc.derivs = FALSE,
                                                                                                                                            optCtrl = list(method = "nlminb", starttests = FALSE, kkt = FALSE)))
anova(interactive,interactive1, test='chisq')

# HO

hinteractive=glmer(human_accuracy~tilt1*clarity1*distadj + (1|video/frame/track_id), family=binomial, data=data2, control = glmerControl(optimizer = "optimx", calc.derivs = FALSE,
                                                                                                                                         optCtrl = list(method = "nlminb", starttests = FALSE, kkt = FALSE)))
hinteractive1=glmer(human_accuracy~tilt1+clarity1+distadj + (1|video/frame/track_id), family=binomial, data=data2, control = glmerControl(optimizer = "optimx", calc.derivs = FALSE,
                                                                                                                                          optCtrl = list(method = "nlminb", starttests = FALSE, kkt = FALSE)))
anova(hinteractive,hinteractive1, test='chisq')

### this is to see if ant walkind speed differs across different categories of blurriness
### NB the following model is valid for both BACH and HO since it is just a measure of blurriness 
### also, no need for nested random factors as we do not consider identification. We only implement video as random factor

blur=glmer(distadj10~clarity1 + (1|video) + (1|obs_effect), family=poisson, data=data2)
blur1=glmer(distadj10~ (1|video) + (1|obs_effect) , family=poisson, data=data2)
anova(blur,blur1, test='Chisq')
emm = emmeans(blur, ~clarity1) #here I create the contrasts 
pairs(emm) #here I test the contrasts


###interaction tilt and clarity

# BACH
tiltclar=glmer(computer_accuracy~tilt1*clarity1 + (1|video/frame/track_id), family='binomial', data=data1, control = glmerControl(optimizer = "optimx", calc.derivs = FALSE,
                                                                                                                                  optCtrl = list(method = "nlminb", starttests = FALSE, kkt = FALSE)))
tiltclar1=glmer(computer_accuracy~tilt1+clarity1 + (1|video/frame/track_id), family='binomial', data=data1, control = glmerControl(optimizer = "optimx", calc.derivs = FALSE,
                                                                                                                                   optCtrl = list(method = "nlminb", starttests = FALSE, kkt = FALSE)))
anova(tiltclar,tiltclar1, test='chisq')
emm = emmeans(tiltclar, ~ clarity1) #here I create the contrasts 
pairs(emm) #here I test the contrasts

# here for separate conditions, but nest does not work because all images are focused
# BACH

tiltclarnest=glmer(computer_accuracy~tilt1+ (1|video/frame/track_id), family='binomial', data=subset(data1, condition=='nestinfrared'), control = glmerControl(optimizer = "optimx", calc.derivs = FALSE,
                                                                                                                                                               optCtrl = list(method = "nlminb", starttests = FALSE, kkt = FALSE)))
tiltclarnestx=glmer(computer_accuracy~ (1|video/frame/track_id), family='binomial', data=subset(data1, condition=='nestinfrared'), control = glmerControl(optimizer = "optimx", calc.derivs = FALSE,
                                                                                                                                                          optCtrl = list(method = "nlminb", starttests = FALSE, kkt = FALSE)))
anova(tiltclarnest,tiltclarnestx, test='Chisq')
emm = emmeans(tiltclarnest, ~ tilt1) #here I create the contrasts 
pairs(emm) #here I test the contrasts

### now foraging area

tiltclarFI=glmer(computer_accuracy~tilt1*clarity1 + (1|video/frame/track_id), family='binomial', data=subset(data1, condition=='foraginginfrared'), control = glmerControl(optimizer = "optimx", calc.derivs = FALSE,
                                                                                                                                                                           optCtrl = list(method = "nlminb", starttests = FALSE, kkt = FALSE)))
tiltclarFIx=glmer(computer_accuracy~tilt1+clarity1 + (1|video/frame/track_id), family='binomial', data=subset(data1, condition=='foraginginfrared'), control = glmerControl(optimizer = "optimx", calc.derivs = FALSE,
                                                                                                                                                                            optCtrl = list(method = "nlminb", starttests = FALSE, kkt = FALSE)))
anova(tiltclarFI,tiltclarFIx, test='Chisq')
emm = emmeans(tiltclarFI, ~ tilt1*clarity1) #here I create the contrasts 
pairs(emm) #here I test the contrasts


tiltclarFV=glmer(computer_accuracy~tilt1*clarity1 + (1|video/frame/track_id), family='binomial', data=subset(data1, condition=='foragingvisible'), control = glmerControl(optimizer = "optimx", calc.derivs = FALSE,
                                                                                                                                                                          optCtrl = list(method = "nlminb", starttests = FALSE, kkt = FALSE)))
tiltclarFVx=glmer(computer_accuracy~tilt1+clarity1 + (1|video/frame/track_id), family='binomial', data=subset(data1, condition=='foragingvisible'), control = glmerControl(optimizer = "optimx", calc.derivs = FALSE,
                                                                                                                                                                           optCtrl = list(method = "nlminb", starttests = FALSE, kkt = FALSE)))
anova(tiltclarFV,tiltclarFVx, test='Chisq')
emm = emmeans(tiltclarFV, ~ tilt1*clarity1) #here I create the contrasts 
pairs(emm) #here I test the contrasts

# HO
tiltclarh=glmer(human_accuracy~tilt1*clarity1 + (1|video/frame/track_id), family='binomial', data=data1, control = glmerControl(optimizer = "optimx", calc.derivs = FALSE,
                                                                                                                                optCtrl = list(method = "bobyqa", starttests = FALSE, kkt = FALSE)))
tiltclar1h=glmer(human_accuracy~tilt1+clarity1 + (1|video/frame/track_id), family='binomial', data=data1, control = glmerControl(optimizer = "optimx", calc.derivs = FALSE,
                                                                                                                                 optCtrl = list(method = "bobyqa", starttests = FALSE, kkt = FALSE)))
anova(tiltclarh,tiltclar1h, test='chisq')
emm = emmeans(tiltclarh, ~tilt1*clarity1) #here I create the contrasts 
pairs(emm) #here I test the contrasts

#-----------

tiltclarnesth=glmer(human_accuracy~tilt1+ (1|video/frame/track_id), family='binomial', data=subset(data1, condition=='nestinfrared'), control = glmerControl(optimizer = "optimx", calc.derivs = FALSE,
                                                                                                                                                             optCtrl = list(method = "nlminb", starttests = FALSE, kkt = FALSE)))
tiltclarnesthx=glmer(human_accuracy~ (1|video/frame/track_id), family='binomial', data=subset(data1, condition=='nestinfrared'), control = glmerControl(optimizer = "optimx", calc.derivs = FALSE,
                                                                                                                                                        optCtrl = list(method = "nlminb", starttests = FALSE, kkt = FALSE)))
anova(tiltclarnesth,tiltclarnesthx, test='Chisq')

emm = emmeans(tiltclarnesth, ~ tilt1) #here I create the contrasts 
pairs(emm) #here I test the contrasts

#-----------

tiltclarFIh=glmer(human_accuracy~tilt1*clarity1 + (1|video/frame/track_id), family='binomial', data=subset(data1, condition=='foraginginfrared'), control = glmerControl(optimizer = "optimx", calc.derivs = FALSE,
                                                                                                                                                                         optCtrl = list(method = "bobyqa", starttests = FALSE, kkt = FALSE)))
tiltclarFIxh=glmer(human_accuracy~tilt1+clarity1 + (1|video/frame/track_id), family='binomial', data=subset(data1, condition=='foraginginfrared'), control = glmerControl(optimizer = "optimx", calc.derivs = FALSE,
                                                                                                                                                                          optCtrl = list(method = "bobyqa", starttests = FALSE, kkt = FALSE)))
anova(tiltclarFIh,tiltclarFIxh, test='Chisq')
emm = emmeans(tiltclarFIh, ~ tilt1*clarity1) #here I create the contrasts 
pairs(emm) #here I test the contrasts

#-----------

tiltclarFVh=glmer(human_accuracy~tilt1*clarity1 + (1|video/frame/track_id), family='binomial', data=subset(data1, condition=='foragingvisible'), control = glmerControl(optimizer = "optimx", calc.derivs = FALSE,
                                                                                                                                                                        optCtrl = list(method = "bobyqa", starttests = FALSE, kkt = FALSE)))
tiltclarFVxh=glmer(human_accuracy~tilt1+clarity1 + (1|video/frame/track_id), family='binomial', data=subset(data1, condition=='foragingvisible'), control = glmerControl(optimizer = "optimx", calc.derivs = FALSE,
                                                                                                                                                                         optCtrl = list(method = "bobyqa", starttests = FALSE, kkt = FALSE)))
anova(tiltclarFVh,tiltclarFVxh, test='Chisq')
emm = emmeans(tiltclarFVh, ~ tilt1*clarity1) #here I create the contrasts 
pairs(emm) #here I test the contrasts


#to calculate percentages of identifications and correct identifications
#HO
table(data1$human.code)
table(data1$human_accuracy)

table(subset(data1, condition=='nestinfrared')$human.code)
table(subset(data1, condition=='nestinfrared')$human_accuracy)

table(subset(data1, condition=='foraginginfrared')$human.code)
table(subset(data1, condition=='foraginginfrared')$human_accuracy)

table(subset(data1, condition=='foragingvisible')$human.code)
table(subset(data1, condition=='foragingvisible')$human_accuracy)

#BACH

table(data1$computer_code)
table(data1$computer_accuracy)

table(subset(data1, condition=='nestinfrared')$computer_code)
table(subset(data1, condition=='nestinfrared')$computer_accuracy)

table(subset(data1, condition=='foraginginfrared')$computer_code)
table(subset(data1, condition=='foraginginfrared')$computer_accuracy)

table(subset(data1, condition=='foragingvisible')$computer_code)
table(subset(data1, condition=='foragingvisible')$computer_accuracy)

#chisquares to test whether correct detections to detections ratios differ between identifiers across conditions

nest = matrix(c(3,49,963,814),ncol=2)
chisq.test(nest, correct=F)

FI = matrix(c(10,16,598,284),ncol=2)
chisq.test(FI, correct=F)

FV = matrix(c(0,61,796,365),ncol=2)
chisq.test(FV, correct=F)
```
